# Isolation and characterisation of novel phages infecting *Lactobacillus plantarum* and proposal of a new genus, “Silenusvirus”

**DOI:** 10.1101/728261

**Authors:** Ifigeneia Kyrkou, Alexander Byth Carstens, Lea Ellegaard-Jensen, Witold Kot, Athanasios Zervas, Amaru Miranda Djurhuus, Horst Neve, Charles M.A.P. Franz, Martin Hansen, Lars Hestbjerg Hansen

## Abstract

Bacteria of *Lactobacillus* sp. are very useful to humans. However, the biology and genomic diversity of their (bacterio)phage enemies remains understudied. Knowledge on *Lactobacillus* phage diversity should broaden to develop efficient phage control strategies. To this end, organic waste samples were screened for phages against two wine-related *Lactobacillus plantarum* strains. Isolates were shotgun sequenced and compared against the phage database and each other by phylogenetics and comparative genomics. The new isolates had only three distant relatives from the database but displayed a high overall degree of genomic similarity amongst them. The latter allowed for the use of one isolate as a representative to conduct transmission electron microscopy and structural protein sequencing, and to study phage adsorption and growth kinetics. The microscopy and proteomics tests confirmed the observed diversity of the new isolates and supported their classification to the family *Siphoviridae* and the proposal of the new phage genus “Silenusvirus”.

## Introduction

Lactic acid bacteria are microorganisms of profound value to humans. They protect food and feed products from spoilage bacteria via acidification and act as sensory biomodulators by fermenting different food matrices^1^. Moreover, some benefits of those lactic acid bacteria recognised as probiotics entail health-promoting effects^2^. Lactic acid bacteria are encompassed within the genera of *Lactococcus, Pediococcus, Streptococcus, Enterococcus, Oenococcus, Leuconostoc, Lactobacillus, Fructobacillus, Weissella* and *Carnobacterium*^3–5^.A member of the genus *Lactobacillus, L. plantarum* is a versatile lactic acid bacterium of great potential for the food industry. This is due to the increasing popularity of *L. plantarum* as starter and adjunct culture from dairy to wine fermentations^6, 7^. Not least, many studies have suggested that supplementation with this bacterial species can have many advantages to the health and welfare of crop plants^8^. Nevertheless, the various applications of *L. plantarum* are at stake because of possible disruptions by bacteriophages (phages) infecting these bacteria, as well reported for other lactic acid bacteria^8–10^ and probiotic strains of *L. plantarum*^6^.

Effective control strategies against phages of *Lactobacillus* sp. can be facilitated with deep knowledge of their biology and genomic diversity. Unfortunately, the diversity of most reported *Lactobacillus* phages has not been thoroughly addressed^11^. At the same time, the current taxonomy of *Lactobacillus* phages has been limited to two families, *Herelleviridae* and *Siphoviridae*, although a proposal that extends it to the family *Myoviridae* has been recently published^12^.The *Herelleviridae* family hosts just two species members, *Lactobacillus virus Lb338-1* and *Lactobacillus virus LP65*, while *Siphoviridae* includes the two genera of *Lactobacillus* phages to have been officially approved by the International Committee on Taxonomy of Viruses (ICTV)^13^. The first genus is called “*Cequinquevirus*” and contains the species *Lactobacillus virus C5, Ld3, Ld17, Ld25A, LLKu* and *phiLdb*^14, 15^. The second genus is called “*Coetzeevirus*” and involves the species *Lactobacillus virus phiJL-1, Pediococcus virus clP1* and *Lactobacillus virus ATCC 8014-B1*^15^. The objective of this study was to provide more insight into the diversity of *Lactobacillus* phages by examining for the first time a group of newly isolated phages that target the industrially relevant *L. plantarum*.

## Methods

### Environmental samples, phage assays and bacterial strains

*Lactobacillus* phages were isolated from organic household waste samples. The samples were collected from two different organic waste treatment plants in Denmark (treatment plants A and B). Initially, the samples were split into two subsamples and processed as detailed in Kyrkou *et al*^12^. The resulting filtrates were screened for phages using the double agar overlay method^16^ and a top layer of 0.4% w/v agarose. Specifically, efficiency of plating assays were performed against two indicator strains that had earlier been acquired from private collections, *L. plantarum* L1 (wine fermentation isolate) and *L. plantarum* MW-1 (grape isolate). Single plaques were resuspended in 0.7 mL SM buffer and later filtered by 0.45-µm pore size PVDF spin filters (Ciro, Florida, USA). Each purified plaque underwent two further reisolation-filtration cycles to ensure pure stock cultures and was stored at 4 °C. For transmission electron microscopy (TEM) and protein sequencing, lysates of 10^10^ plaque-forming units (PFUs)/mL) were further purified and concentrated using caesium chloride (CsCl) gradient ultracentrifugation, as described elsewhere^17^. All incubations of phage manipulations were done at 25 °C using De Man, Rogosa and Sharpe (MRS) broth and agar media supplemented with 10 mM CaCl_2_ (MRSΦ), whereas indicator strains were grown in MRS at 37 °C.

### TEM analysis and structural protein sequencing

Phage morphology and structural proteins were determined using the CsCl-purified stocks. Micrographs of phage Silenus were generated as in other studies^12, 18^. The mean values and standard deviations of all Silenus virion dimensions were elucidated after inspection of 20-23 phage particles. Structural proteins were sequenced following published protocols^12, 18^. Briefly, 100 µL of the CsCl-purified stocks were filtered through an Amicon Ultra filter unit (MWCO 30k Da) and desalted four times. Phage particles (10 µL) were denaturised in 6 M urea, 5 mM dithiothreitol and 50 mM Tris-HCl (pH 8) and destabilised by freeze-thawing. Proteins were reduced (1h incubation, 60 °C) and alkylated in 100 mM iodoacetamide and 50 mM ammonium bicarbonate, digested with 0.8 µg trypsin in 50 mM ammonia bicarbonate (40 µL) and diluted in 0.05% trifluoroacetic acid. The resulting peptides were analysed with an Ultimate 3,000 RSLCnano UHPLC system coupled with an analytical column (75 µm × 250 mm, 2 µm C18) and a Q Exactive HF mass spectrometer (ThermoFisher Scientific, Denmark). The twelve most intense ions were selected using MS Orbitrap scans and subsequently MS/MS-fragmented at a normalised collision energy (28) and a resolution of 60,000 (m/z 200). The output data were analysed in Proteome Discoverer 2.2 (ThermoFisher Scientific) and searched against predicted phage proteins by the Sequest HT algorithm.

### Phage DNA isolation, library construction and sequencing

Phage DNA was extracted from the filtered stock lysates according to a standard phenol/chloroform method^19^. Sequencing libraries were constructed with the Nextera® XT DNA kit (Illumina Inc., San Diego, California, USA) according to the manufacturer’s instructions for library preparations. The library normalisation, pooling and sequencing were done as indicated elsewhere^20^. All phage genomes were sequenced as a part of a flowcell on the Illumina MiSeq platform using the v2, 2×250 cycles chemistry.

### Bioinformatics analyses

*De novo* genome assembling was done with SPAdes (v. 3.5.0)^21^ and the assemblies were cross-verified with Unicycler (v. 0.4.3)^22^ and CLC Genomic Workbench (v. 9.5.3; CLC bio, Aarhus, Denmark) according to already published methods^12, 23^. The annotation pipeline involved automatic protein annotations with RASTtk^24^ and GeneMark^25^ as a gene caller, followed by manual curations based on the predictions of BLASTp, HHpred^26^ and sometimes PfamScan^27^. Lysin amino acid sequences were multiply aligned with Clustal Omega and viewed with MView (v. 1.63) using the default settings^28^. Transmembrane domains were identified with TMHMM^29^. The genome of phage Sabazios was scanned for −1 frameshift slippery sequences near the two lysin genes with FSFinder^30^ and all genomes were scanned for tRNA genes with tRNAscan-SE (v. 2.0)^31^.

For each pair of compared query-subject phage genomes, query cover was multiplied by identity, according to the crude method for estimating nucleotide similarity of ICTV^32^. For more stringent comparisons on the nucleotide level the tool Gegenees^33^ was used, after customising the fragment size/sliding step size (50/25) and threshold (0%). Phylogenetic analyses were conducted for phages Silenus, Sabazios, Bassarid, and compared to phages that returned a Gegenees score of at least 0.05. Two phylogenetic trees were built with the default pipeline of “One Click mode” (http://phylogeny.Lirmm.fr/)^34^. The first tree was based on the major capsid protein, the second on the large subunit of terminase. All-against-all protein homology checks were performed between and within proteomes of the phages studied here and their closest relatives. The CMG-biotools package^35^, which implements the BLASTp algorithm, was chosen for this purpose. Pairs of proteins that aligned for at least 50% of the longest sequence and shared at least 50% of their nucleotides within the aligned region were considered as positive hits. Visualisation of genome alignments among the phages of this study and their two closest phage relatives were done with Easyfig^36^ and the BLASTn algorithm.

### Phage adsorption and growth kinetics

Phage adsorption and one-step growth experiments were done at a multiplicity of infection of 0.05. Strain MW-1 was grown to an OD_600_ of 3.2, which corresponds to approximately 10^8^ colony-forming units (CFUs)/mL for this strain. Immediately after, MW-1 cultures were infected with phage Silenus and incubated for 10 min at 37 °C. This time of infection was recorded as time point zero. The remaining steps of the adsorption assay and the burst size assay followed an already published protocol^18^ under shaking conditions and at an incubation temperature of 37 °C. Phage growth kinetics were monitored on four-fold dilutions of the infected MW-1 cultures in triplicate assays. Samples were harvested from each triplicate assay approximately every 5-10 min, serially diluted and plated against a lawn of MW-1, incubated overnight and then examined for plaques. Supernatants of the infected MW-1 cultures just before the dilution step were plated, as well. The total count of PFUs (unadsorbed phages) from these supernatants designated how many plaques should be disregarded when counting infected centers.

### Genomic data availability

Assembled and annotated genomes of phages Silenus, Bassarid and Sabazios were uploaded to GenBank under accession numbers MG765278, MG765275 and MH809528, respectively.

## Results and Discussion

### Phage isolation and burst size

Phages Silenus, Sabazios and Bassarid were all isolated from the organic waste samples. Phages Silenus and Sabazios came from treatment plant B and phage Bassarid from treatment plant A. The phages formed plaques of approximately 1 mm following an incubation period of 24 h at 25 °C in MRSΦ broth and agar media (see Supplementary Fig. S1 online). Lactobacillus phage Bassarid was isolated after infecting the lawn of *L. plantarum* L1, while the other two phages were active against *L. plantarum* MW-1. Adsorption and one-step growth curve tests were performed using Silenus as the representative of the three phages, due to the overall high genomic and morphological similarities among the three phages (details in “Phage morphology”, “Support for the description of a new *Lactobacillus* phage genus” below). The adsorption rate for Silenus was 99.7%, the latent period was 45±5 min and the average burst size was 4.86 progeny virions. Although the burst size of Silenus was low, similar burst sizes have been reported for other *Lactobacillus* phages^37–39^.

### Phage morphology

Most virion-associated genes among phages Silenus, Sabazios and Bassarid were highly conserved (please read “Support for the description of a new *Lactobacillus* phage genus” and Fig. 5 there) and their virion morphologies were similar (see Supplementary Table S1 and Fig. S2 online). For these reasons, phage Silenus was selected as the representative of the three phages for the morphology-related analyses. Transmission electron micrographs displayed phage particles with an isometric head (diameter: 55.6 ± 3.2 nm), a neck passage with thin whiskers, a non-contractile flexible tail (length counting in the baseplate: 173.6 ± 5.6 nm; width: 12.0 ± 0.5 nm) and a characteristic double-disc baseplate (length: 12.2 ± 1.0 nm; width: 20.2 ± 1.4 nm) which culminates in short flexible appendages with tiny terminal globular structures. Similar flexible globular structures and capsid-tail dimensions have been observed for other *Lactobacillus* phages but their exact role remains unknown^3, 40^. Interestingly, in some *Leuconostoc mesenteroides* phages similar baseplate appendages could also adsorb to other parts of the tail in an inconsistent manner^41^. These characteristics classify Silenus to the order *Caudovirales* and the family *Siphoviridae* (Fig. 1), which is the most widespread taxonomic classification among *Lactobacillus* phages^40^.

**Figure 1.**
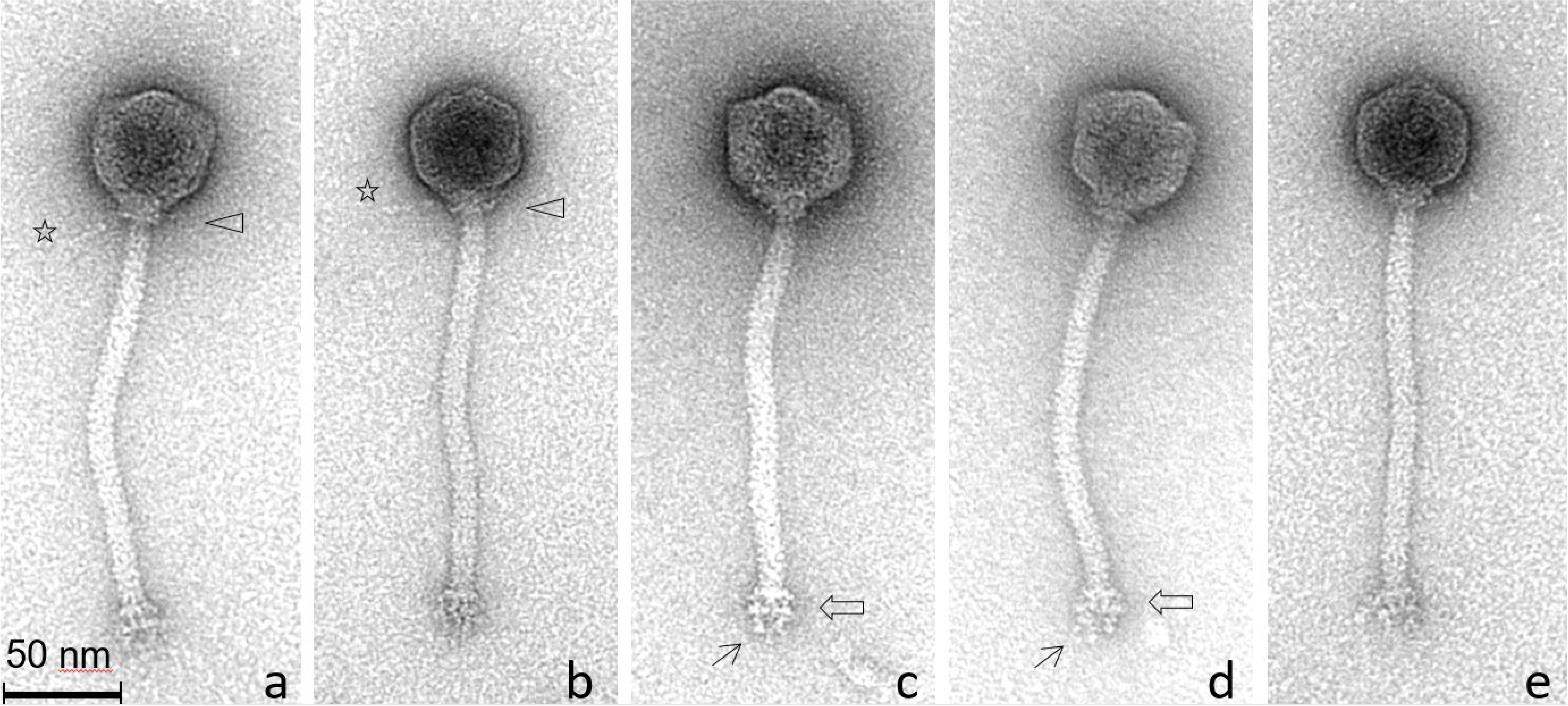
Transmission electron micrographs of *L. plantarum* phage Silenus. In **a** and **b**, triangles indicate the neck passage structure with (faint) whisker structures (see stars). Open arrows in **c** and **d** highlight the characteristic double-disc baseplate structure at the distal end of the flexible, non-contractile tail. Single arrows in **b** and **c** show representative short flexible appendages (with tiny terminal globular structures) attached under the baseplate structures. The observed morphology classifies phage Silenus to the family *Siphoviridae*.

**Figure 2.**
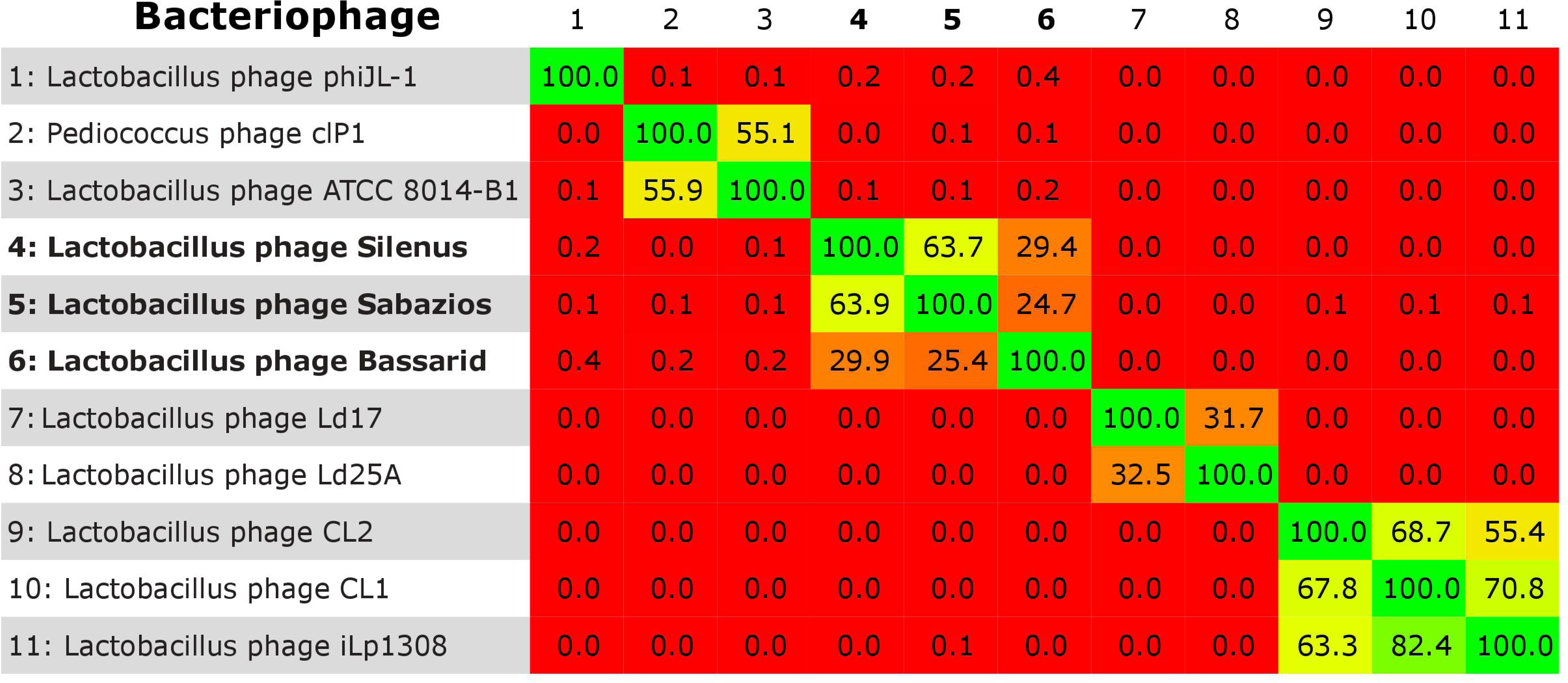
BLASTn heatplot of Gegenees. Red areas illustrate phage pairs with no similarity. The new phages (numbers **4-6**) form a separate group.

**Figure 3.**
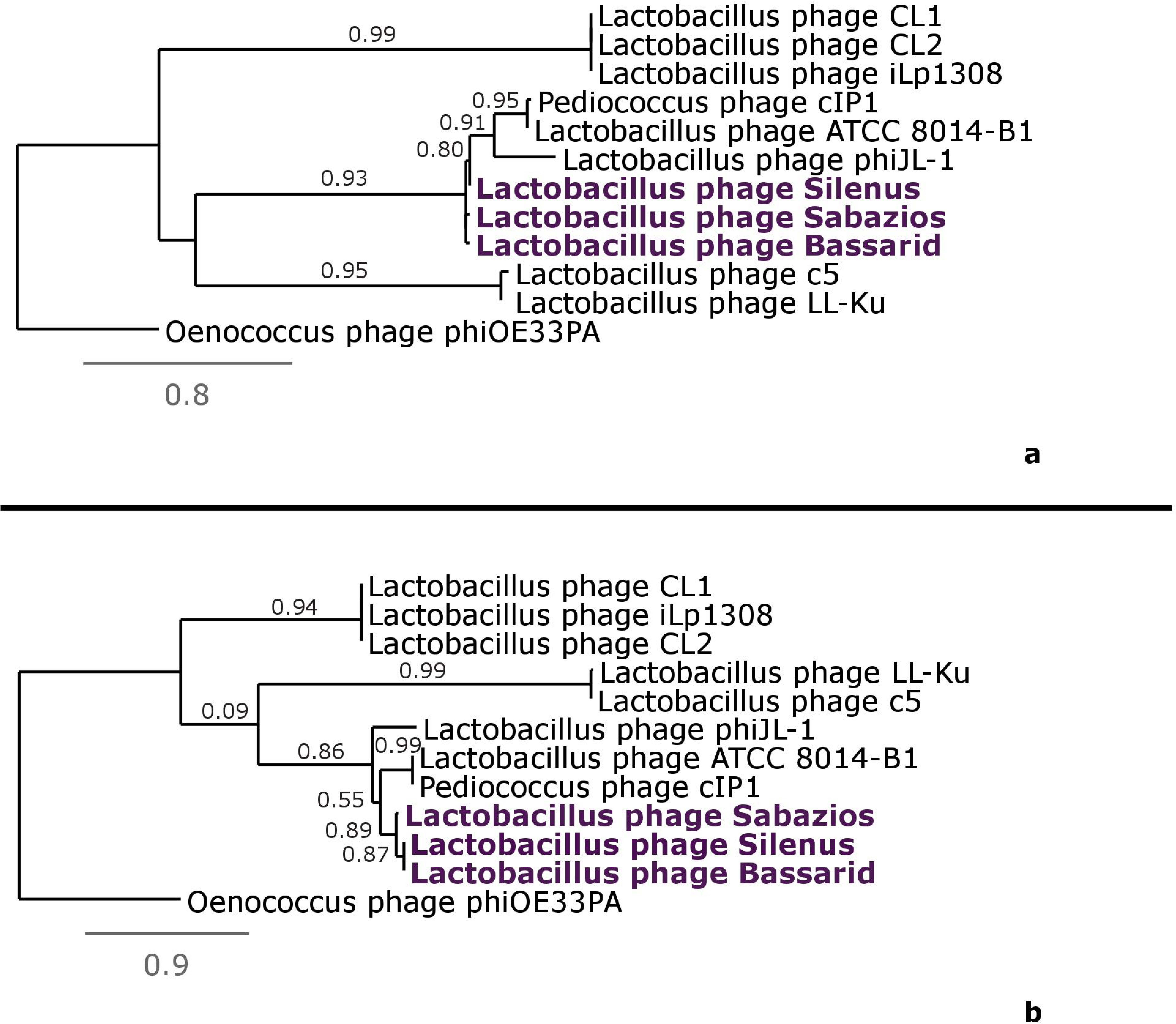
Phylogenetic trees constructed for phages Silenus, Bassarid, Sabazios (given in purple-coloured boldface) and those *Lactobacillus* phages that scored an average similarity of at least 0.05 or higher with Gegenees. Tree **a** was constructed using the amino acid sequences of the major capsid protein. Tree **b** was constructed using the amino acid sequences of the large subunit terminase. Comparisons were run with the “One Click mode” (http://phylogeny.Lirmm.fr/) and Oenococcus phage phiOE33PA proteins as an outgroup.

**Figure 4.**
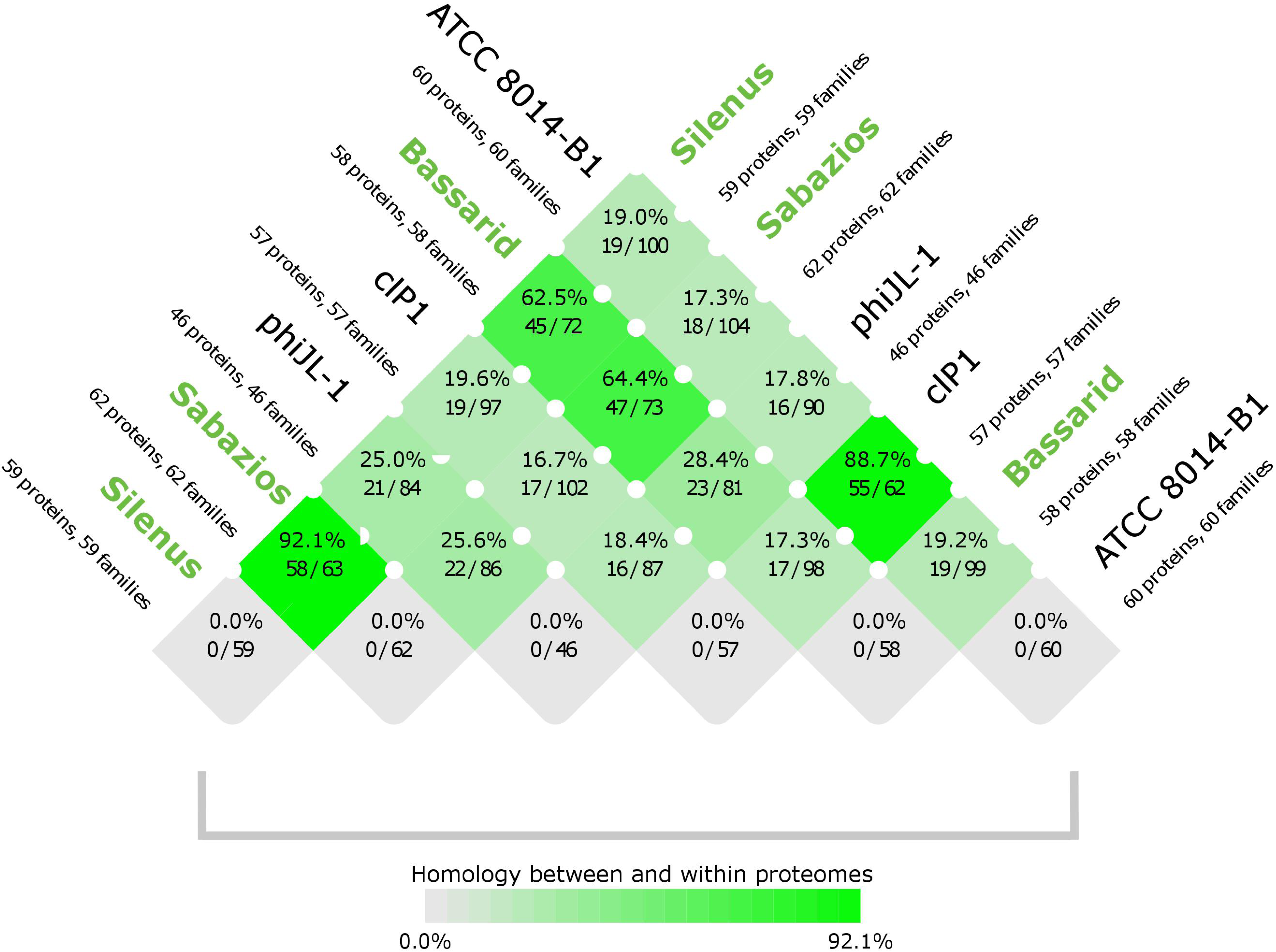
Homology scores between proteomes and within proteomes (last row) of the new phages (given in green boldface) and their closest phage relatives. The intense green colours signify related phage pairs with high (>50%) proteome homology. The faded green to grey colours signify unrelated phage pairs with low (<50%) proteome homology. The scoring was performed with CMG-biotools system.

**Figure 5.**
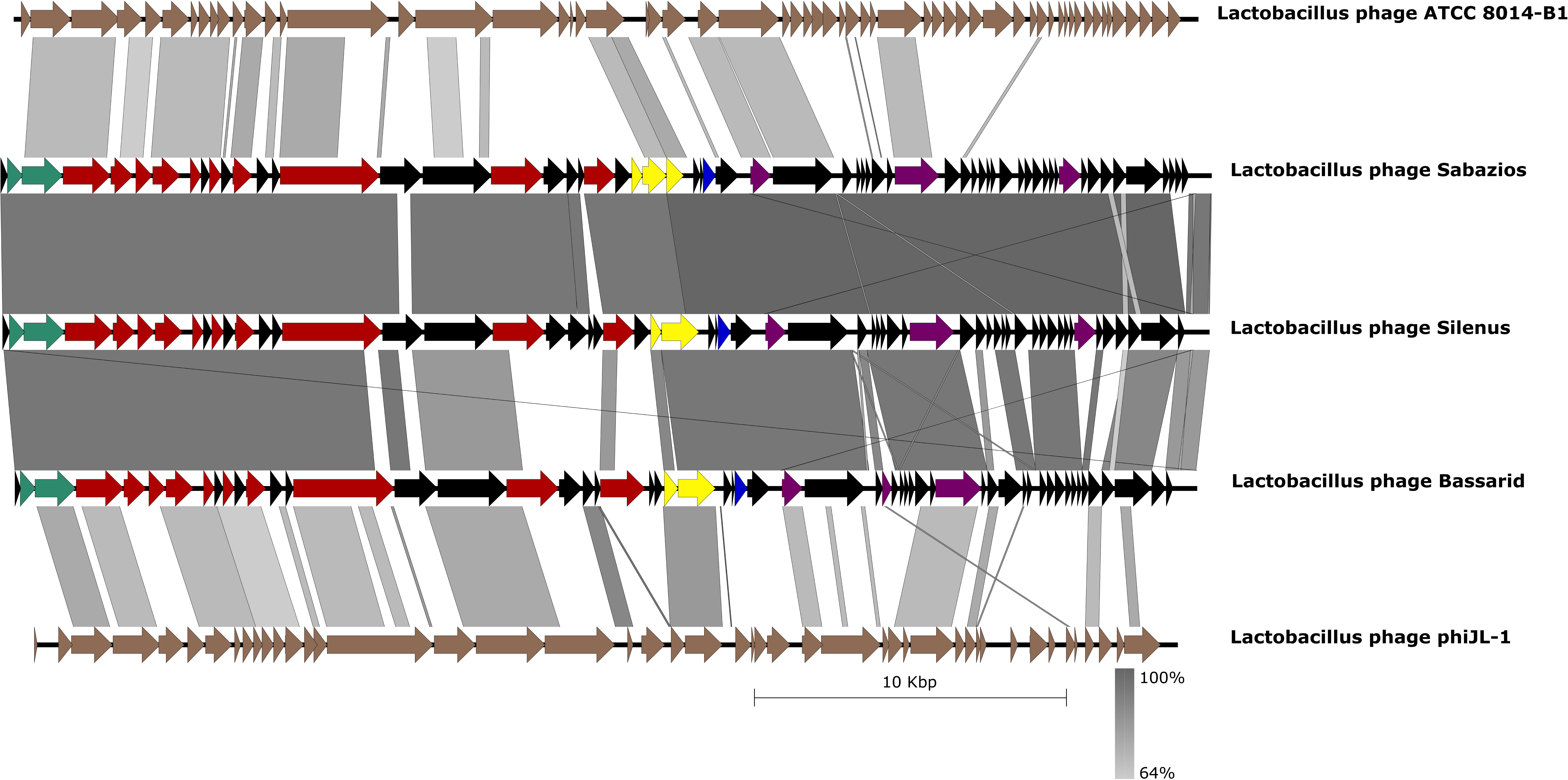
Genomic synteny comparisons with Easyfig and the BLASTn algorithm. The genomes of the new phages are tandemly compared to each other and to their distantly related phages ATCC 8014-B1 and phiJL-1. Arrows represent the locations of coding sequences and shaded lines reflect the degree of homology between pairs of phages. Colours other than black mark specific predicted protein functions; DNA packaging is in turquoise, morphogenesis in red, lysis in yellow, selfish genetic elements in blue and metabolism/modification of nucleic acids in deep purple.

### Basic genomic characteristics

Sequencing reads for each phage were assembled into single contigs of high coverage (667.7x-2,063x). The genome statistics of Silenus, Sabazios and Bassarid are presented in Table 1. Their genome sizes render them some of the smallest phages infecting *L. plantarum* together with phage phiJL-1 and phage ATCC 8014-B1 (accession numbers: NC_006936 and NC_019916), while their G/C content (42.5%) is close to that of their host (44.5%). In the three 37.9-38.8 kbp, double-stranded (ds) DNA phage genomes, all predicted open reading frames (ORFs) were located at the sense strand, which is in accordance with other phages of *L. plantarum*^42^ and the closely-related *P. damnosus*^43^.The total number of predicted ORFs was 58-62 for the three phages. Out of this total, specific functions were assigned to 18 coding sequences for phages Bassarid and Silenus and to 19 for phage Sabazios. This translated to a mean of 30.7% of their coding sequences based on the in silico open reading frame predictions (Table 1).

**Table 1.**
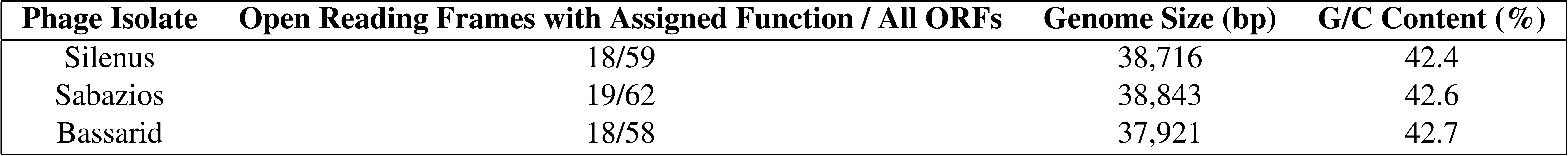
Overall genome statistics of the three *Lactobacillus* phage isolates.

### Description of phage modules

Three different genetic modules could be discerned, which enable the following: DNA packaging, morphogenesis and lysis (see Supplementary Fig. S3-S5 online). On the whole, such a pattern of genome organisation corresponds to most *Lactobacillus* phage isolates^3^. Most of the proteins that could be assigned functions to were identified in all three phages.

#### DNA packaging module proteins

The DNA packaging modules of Silenus, Sabazios and Bassarid contained genes that encode a small subunit terminase and a large subunit terminase. Downstream of the terminase unit, the sequencing coverage increased nearly twofold for all three phages. This may indicate a *pac*-type headful packaging strategy, where the variable cleaving sites for packaging termination lead to some regions being dual in some phage particles^44^. A *pac*-type headful packaging was experimentally determined for the distant relatives of the three new phages, Lactobacillus phage ATCC 8014-B1 and Lactobacillus phage phiJL-1^42, 45^. For these reasons, the start of the new phages’ genomes was arbitrarily set to the coding sequence of a hypothetical protein just before the small subunit terminase. In double-stranded DNA phages, the small subunit terminase is responsible for the specific recognition and binding to the *cos* or *pac* sequence of the phage DNA^46^. The main functions of the large subunit are three; the large subunit supplies the packaging motor with the ATPase activity needed for DNA translocation, it cleaves DNA concatemers during prohead filling and it interacts with the prohead portal protein of the vertex^46^. Studies on the phylogeny of large subunit terminases have proven that these are descendants of a common ancestor, hence they are conserved among related phages^47^. Consequently, the large subunit terminase was chosen to construct a phylogenetic tree of the phages of this study (Fig. 3b). The terminase unit of the new phages was located next to the morphogenesis module, as typically seen in other phages^46^. In all three genomes, the portal protein constituted the junction of the DNA packaging and the morphogenesis module. Indeed, the portal protein is a crucial component of the packaging motor, because it appears to pump the phage genome into the capsid with the aid of the large subunit terminase (ATPase activity)^48^. Moreover, it has been proposed that the portal protein directs the final shape and size of the prohead^49, 50^. An HHpred search revealed that the extended packaging (i.e. including the portal protein) region of the three new phages was highly conserved (Fig. 5). Altogether, the interrelated *Bacillus* phages SF6 and SPP1 matched that region better than phages ATCC 8014-B1 and phiJL-1. This finding supports the *pac*-type nature of the new phages, since phages SF6 and SPP1 were shown to rely on *pac* packaging^51, 52^.

#### Morphogenesis module proteins

Predicted proteins of known function within the morphogenesis module of the three phages included proteins of the capsid (a scaffolding protein and a major capsid protein), proteins of the connector (the portal protein and a putative head-to-tail joining protein) and proteins of the tail (a major tail protein, a putative tape measure protein and a tail fiber protein). LC-MS/MS analysis verified that the representative phage Silenus produces three of the aforementioned proteins, one protein of the capsid (major capsid), one of the connector (putative head-to-tail joining protein) and a hypothetical protein (peg. 5; see Supplementary Table S2 online). The latter probably belongs to the tail, since its gene is located right downstream of the putative tape measure gene. Two more proteins have been sequenced but whether these constitute virion-associated proteins or the result of contamination is unclear (peg. 27 and peg. 48; Supplementary Table S2). As expected, the scaffolding protein was not traced by the protein sequencing analysis, because it comprises the core of the pre-assembled prohead, which is surrounded by the major capsid protein^53^. The scaffolding protein is absent from mature capsids, since it is removed to make room for the phage genome, and its removal triggers a reaction that stabilises the structure of the capsid^54, 55^.The putative head-to-tail joining protein of the phages of this study can belong either to head-completion proteins, i.e. adaptors and stoppers, or to a tail-completion protein. The proteins of the capsid and the putative head-to-tail joining protein were quite conserved among the new phages and largely aligned with their orthologues in *Lactobacillus* phages ATCC 8014-B1 and phiJL-1, and Pediococcus phage cIP1 (BLASTp results). Similarly, the major tail and the putative tape measure proteins of Silenus, Sabazios and Bassarid contained domains with homology to the aforementioned phages and primarily to Lactobacillus phage phiJL-1. One or two different major tail proteins are principal units of the tail tube, while their role varies from phage to phage^56, 57^. The tape measure protein, another principal unit of the tail, is enclosed in a shell of major tail proteins^58^. In the genomes of the new phages, the tape measure protein, whose length defines the length of the tail, had the longest sequence (1,066-1,078 aa) and was located downstream of the major tail protein^59, 60^. The predicted tail fiber protein of the new phages is likely to initiate the infection process through the identification and binding to host receptors on the surface of sensitive cells, as described for other phages^61, 62^. An interesting feature of the new phages’ genomes was that the tail fiber protein and all hypothetical proteins found in the region between the tape measure protein and the lysis module aligned poorly with existing phage records. In average, this region exhibited low nucleotide similarity between phage Bassarid and phages Silenus, Sabazios, as well (BLASTp results; see also Fig. 5). Tail fiber proteins identify host receptors with great specificity. The apparent horizontal transferability of tail fiber genes between phage species reshapes the host range of phages in a constant manner^63, 64^. Considering these, it can be expected that the host range of phage Bassarid varies from that of the other two phages.

#### Lysis module proteins

A classic phage lysis cassette comprises two types of proteins, a holin and a lysin. These proteins are interdependent and both of them are generally essential for the successful release of dsDNA phage progeny^65^. Holins permeabilise the cytoplasmic membrane thereby granting access of the cell wall to lysins, which then degrade the cell wall^66^. In this way, holins virtually control when lysis should occur^65^. Usually, holins have two or three transmembrane domains, but the holins of the phages described in this study belonged to the rare one-transmembrane domain group^67^ (TMHMM search). Additionally, their best match was ORF147 from phage phiJL-1, that is likely a holin with one transmembrane domain^42^. Studies indicate that *Lactobacillus* phages only employ two out of the five classes of lysins, muramidases and amidases^68^. All lysins of the phages studied here strongly matched muramidase entries (HHpred search). In this study, noteworthy was the prediction of two slightly overlapping ORFs for lysins within the module of phage Sabazios. According to Clustal Omega alignments, the first amino acid sequence of Sabazios lysins aligned with high homology to the first 64.8% of Bassarid’s and Sabazios’ lysin sequence. The second amino acid sequence of Sabazios lysins aligned with a small overlap to the first and with high homology to the last 46.5% of Bassarid’s and Sabazios’ lysin sequence (see Supplementary Fig. S6 online). It is possible that these two lysins are products of a nonsense or frameshift mutation, which would create a pair of non-functional pieces^69^. Alternatively, the two overlapping CDS may produce the subunits of a dimeric lysin, like those of phages CD27L and CTP1L of *Clostridium difficile*^70^. A third option would be a programmed ribosomal slippage, which may either adjust the degree of lysin production as a response to conditions in the cell or produce a specific ratio of two different lysin proteins^71^. So far, ribosomal slippage has principally been reported for structural genes of the tail^72^. In the only studied case of slippage for *Lactobacillus* phages, both the major capsid and the major tail protein of phage A2 are affected by ribosomal slippage^73, 74^. However, FSFinder did not trace any slippery sequence in the overlap region of the two lysin ORFs of phage Sabazios. The fact that all predicted ORFs were found at the same strand implies the absence of genetic switches and thus, given that no other lysogeny-related genes were detected, we presume that the three new phages are most probably virulent.

#### Other predicted proteins

The existence of other modules could not be confirmed, but some additional proteins in the genomes of Sabazios, Silenus and Bassarid were assigned to a function. A superfamily II, ATP-dependent helicase, predicted in all three phages, may belong to the family of DEAD/DEAH-box containing helicases (BLASTp search result). Helicases, i.e. enzymes that catalyze the unfolding of DNA or RNA, are often produced by phage genomes^75^. Most notably, DEAD/DEAH-box containing helicases participate in RNA metabolism in many, essential ways, such as by regulating gene expression and signalling^76^. These functions could explain why it is likely for DEAD/DEAH-box containing helicase genes to be found within a phage genome. In all cases, the vast majority of proteins in close proximity to the new phages’ helicase could not be annotated. Along with the helicase, a gene of the phages Sabazios and Silenus encoded an adenine-specific methyltransferase, which is a DNA modification enzyme. A BLASTp search revealed orthologues of Sabazios’ and Silenus’ methyltransferases in the genomes of *L. plantarum* and other *Lactobacillus* sp. This finding corroborates that these two phages mimic the host genome’s methylation pattern as an active strategy to evade restriction by host-driven endonucleases^77^. In phages, methyltransferases are often transferred through horizontal gene transfer. Genes coding for methyltransferases are common in phage genomes where they seem to participate in various functions in addition to nucleic acid methylation^78^. Methyltransferase encoding genes have already been noted in the genomes of some *L. plantarum* strains (REBASE search; http://rebase.neb.com/rebase/rebase.html) and in the genomes of *Lactobacillus* phages PL1, J-1, P1174 (NCBI Protein database search).

Selfish genetic elements in the genomes of Silenus, Bassarid and Sabazios were represented by one HNH homing endonuclease. One or more HNH endonuclease genes have been found in the genomes of *Lactobacillus* phages before^12^. Phage-encoded HNH endonucleases can be part of self-splicing genes, such as group I and II introns and inteins, but they can also be free-standing^79^. Essentially, HNH endonucleases are highly specialised selfish genetic elements that facilitate the mobility of themselves and of those genes to which they pertain, from genome to genome^79^. In some cases, phages that produce HNH endonucleases can even exclude other competing phage species by cleaving their DNA^80^. Due to the position of the new phages’ HNH endonuclease genes next to hypothetical genes, the role of these enzymes could not be deduced. We joined the two ORFs framing each HNH endonuclease of the studied phages and performed a BLASTn analyses. None of the resulting BLASTn hits spanned along the joint region suggesting that no gene was spliced by these HNH endonucleases.

### Support for the description of a new *Lactobacillus* phage genus

Multiplying BLASTn query cover by identity yielded a limited (>50%, ICTV criterion) overall nucleotide similarity of phages Silenus, Bassarid and Sabazios to other phage records. The highest BLASTn (<50%) similarity scores were obtained from Lactobacillus phage phiJL-1, Lactobacillus phage ATCC 8014-B1 and Pediococcus phage cIP1. The three new phages and eight BLASTn genome records that had some level of nucleotide sequence homology to the new phages were afterwards submitted to Gegenees. All-against-all BLASTn comparisons performed according to the Gegenees software resulted in the heatplot of Fig. 2. The scored phylogenomic distances supported a separate grouping of the phages of this study from the other eight phages. Nonetheless, diversity within the group of the new phages was also noted, since phage Bassarid was found to be quite distinct from phages Silenus and Sabazios at the nucleotide level (average normalised similarities of 29.9% and 24.7%).

The phylogenetic trees of the major capsid protein and the large subunit terminase protein corroborated the sorting of the new phages into one individual group (Figs. 3a and 3b, respectively). On the other hand, diversity within the group was reiterated with phage Bassarid showing occasional, yet low variation (Fig. 3b). Consistent with earlier observations was the clustering of the three phages and phages phiJL-1, ATCC 8014-B1 and cIP1 into sister groups. Proteome homology tests were done for the new phages and those phages that appeared to be their distant relatives according to the aforementioned tests (i.e. phiJL-1, ATCC 8014-B1 and cIP1). Homology between and within the six phages further clarified how diverse these phages are from existing phage records. Specifically, the three phages share between 62.5-92.1% of their proteome (Fig. 4), while the homology to the proteomes of phiJL-1, ATCC 8014-B1 and cIP1 is low (<28.4%). Regarding homology within each phage proteome, no paralogous proteins were found for any of the six examined phages (last row of Fig. 4).

Finally, genomic synteny tests with Easyfig visualised the previously manifested homology among the new phages and provided evidence for their conserved genome architecture (Fig. 5). The major differences between phage Bassarid and the other two new phages were localised in the region of the tail-related structural genes, and particularly at those hypothetical ORFs that neighbour the tail fiber gene (see Supplementary Fig. S5 online). Nonetheless, no important structural dissimilarities between the tails of Silenus, Sabazios and the tail of Bassarid were further highlighted by TEM (see Supplementary Table S1 and Fig. S2). At the same time, it is seen from Fig. 4 that even if phages phiJL-1 and ATCC 8014-B1 showed some conserved gene order against the new phages they did score low in nucleotide homology.

Taken together, all these results suggest that *Lactobacillus* phages Silenus, Bassarid and Sabazios form a coherent group and are considerably distinct from all other fully-sequenced phages. Therefore, and in agreement with ICTV criteria^32^, we propose the new *Lactobacillus* phage genus “Silenusvirus”. At present, the new genus should consist exclusively of phage Silenus (founder), Bassarid and Sabazios.

## Conclusions

A few phages of *L. plantarum* have been isolated over the years^40^ but this study reports the first of these which can infect wine-related strains of *L. plantarum*. We characterised three new phage species, Lactobacillus phage Sabazios, Lactobacillus phage Bassarid and Lactobacillus phage Silenus, by means of whole-genome sequencing and *in silico* protein prediction. Furthermore, we assessed the growth parameters of a representative phage and investigated its morphology with TEM. By comparing the phages Sabazios, Bassarid and Silenus to existing phage records, we demonstrated substantial genomic heterogeneity. This heterogeneity was further evident by the fact that we could assign functions to less than one third of the predicted ORFs. Our findings support the creation of the novel *Lactobacillus* phage genus “Silenusvirus”, with phages Silenus, Sabazios and Bassarid as the new members of this proposed genus. The results of this study shed more light on the diversity of *L. plantarum* phages and their hosts, and could aid towards the development of efficient phage control interventions in the future.

## Supporting information

Supplementary files compilation

## Acknowledgements

The authors would like to thank all collaborators of the MicroWine consortium for exchange of ideas and material. We would also like to acknowledge Angela Back from Max Rubner-Institut for technical support in the TEM analysis and the companies BioVækst (treatment plant A) and HCS (treatment plant B) for supplying the waste samples. This research was funded by the Horizon 2020 Program of the European Commission within the Marie Skłodowska-Curie Innovative Training Network “MicroWine” (grant number 643063) and the Danish Research Council for Technology and Production project “Phytoprotect” (grant number DFF – 4184-00070B).

## Author contributions statement

I.K., A.B.C. and L.H.H. conceived the project. I.K., A.B.C., M.H. and W.K. developed the methods. I.K., A.Z. and H.N. performed the analyses. I.K., A.B.C., A.M.D., L.E.J. and H.N. conducted the experiments. I.K., A.B.C., W.K., A.Z., M.H., H.N. and C.F. validated the results. I.K. wrote the first draft and later versions of the paper. I.K., A.Z. and H.N. performed the visualisation of the results. L.E.J. and L.H.H. supervised the project. L.H.H.acquired the grants. All authors reviewed the manuscript.

## Supplementary information

Online.

## Ethics declarations

The authors declare no competing interests.

