## Supplementary files compilation for "Isolation and characterisation of novel phages infecting *Lactobacillus plantarum* and proposal of a new genus, “Silenusvirus”"

### Supplementary information

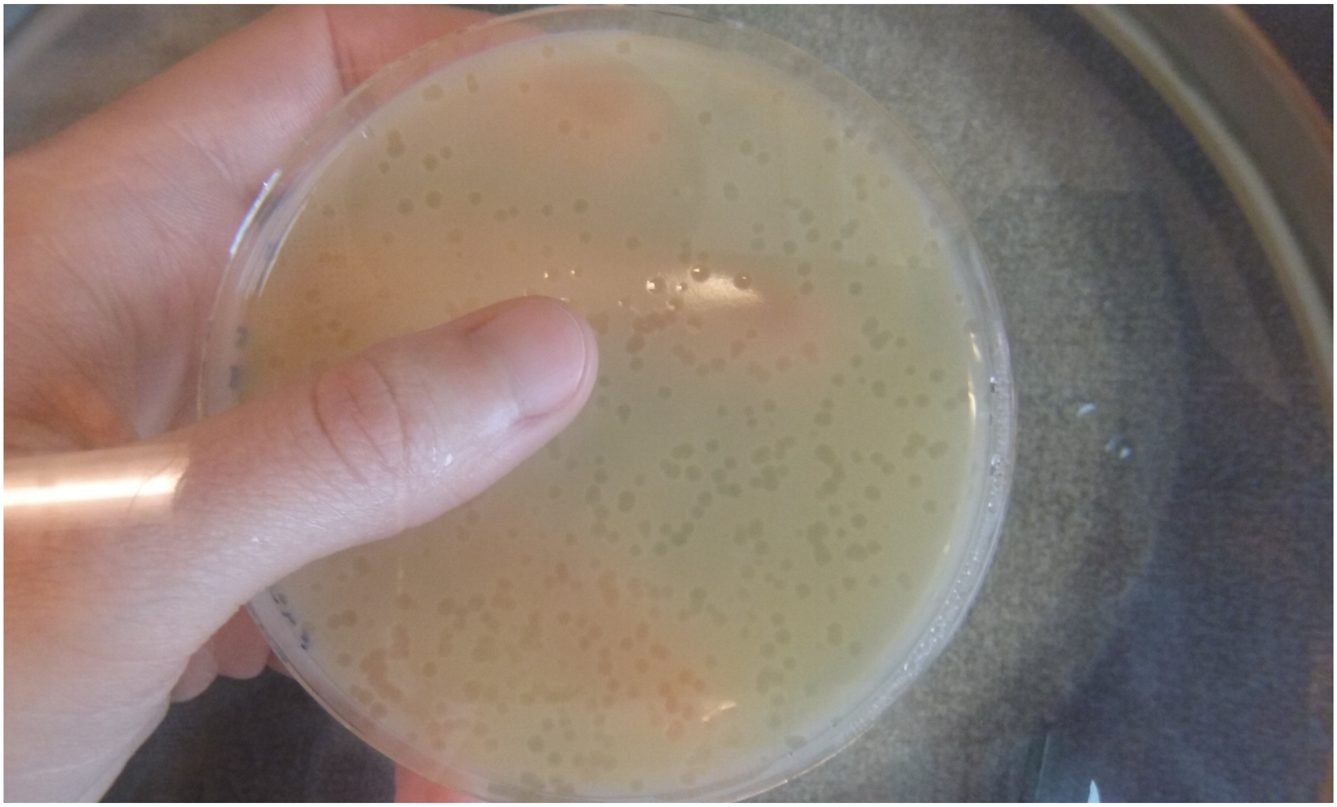

**Figure 1.** Plaques produced on a lawn of *L. plantarum* L1 by phage Bassarid.

| Phage Sabazios | Dimensions (nm) | Counted Phage Particles |
|---|---|---|
| **Head Diameter** | 54.5 ± 2.4 | 18 |
| **Tail Length (with Baseplate)** | 173.1 ± 6.1 | 18 |
| **Tail Width** | 11.5 ± 0.5 | 18 |
| **Baseplate Width** | 17.6 ± 1.3 | 18 |
| **Baseplate Length** | 11.3 ± 1.1 | 18 |
| **Collar Width** | 14.9 ± 0.4 | 3 |

| Phage Bassarid | Dimensions (nm) | Counted Phage Particles |
|---|---|---|
| **(Empty) Head Diameter** | 59.2 ± 1.2 | 4 |
| **Tail Length (with Baseplate)** | 177.0 ± 1.4 | 4 |
| **Tail Width** | 11.7 ± 0.5 | 4 |
| **Baseplate Width** | 18.7 ± 2.2 | 4 |
| **Baseplate Length** | could not be measured | could not be measured |

**Table 1.** TEM analysis results of virion dimensions for phages Sabazios and Bassarid. For the TEM micrographs of these phages see Fig. S1.

| Description | Function | Coverage (%) | No of Peptides | Peptide Spectrum Matches | Molecular Mass (kDa) | Gene |
|---|---|---|---|---|---|---|
| Major capsid protein | Structural | 12 | 3 | 40 | 31.2 | peg. 14 |
| Putative head-to-tail joining protein | Structural | 1 | 1 | 30 | 62.5 | peg. 11 |
| Hypothetical protein | Structural | 3 | 1 | 15 | 49.7 | peg. 5 |
| Adenine-specific methyltransferase | DNA modification | 6 | 1 | 25 | 14.2 | peg. 27 |
| Hypothetical protein | Unknown | 2 | 1 | 1 | 73.1 | peg. 48 |

**Table 2.** Results of the protein sequencing analysis for phage Silenus.

| Bacteriophage record | NCBI nucleotide accession number |
|---|---|
| Lactobacillus phage c5 | NC_019449.1 |
| Lactobacillus phage Ld3 | NC_025421.1 |
| Lactobacillus phage Ld17 | NC_025420.1 |
| Lactobacillus phage Ld25A | NC_025415.1 |
| Lactobacillus phage LL-Ku | NC_022989.1 |
| Lactobacillus phage phiLdb | NC_022762.1 |
| Lactobacillus phage phiJL-1 | NC_006936.1 |
| Lactobacillus phage ATCC 8014-B1 | NC_019916.1 |
| Pediococcus phage clP1 | NC_016161.1 |
| Bacillus phage SPP1 | NC_004166.2 |
| Lactobacillus phage A2 | NC_004112.1 |
| Lactobacillus phage J-1 | NC_022756.1 |
| Lactobacillus phage P1174 | MG913376.1 |
| Lactobacillus phage CL1 | NC_028888.1 |
| Lactobacillus phage CL2 | NC_028835.1 |
| Lactobacillus phage iLp1308 | NC_028911.1 |
| Oenococcus phage phiOE33PA | MH220877.1 |

**Table 3.** NCBI nucleotide accession numbers of all available phage records cited by this article.

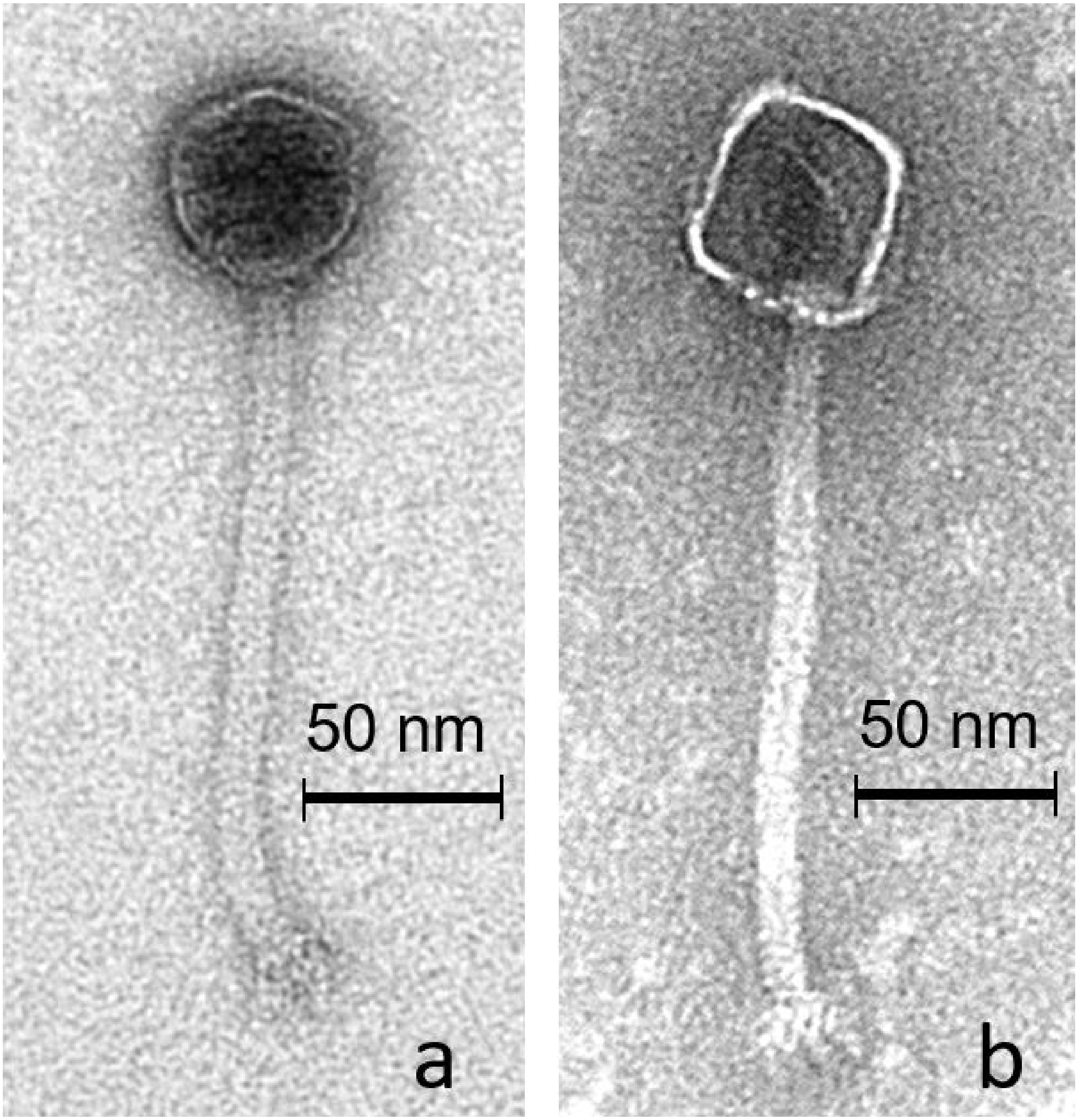


**Figure 2.** TEM micrographs of phages Sabazios (a) and Bassarid (b). For phage Bassarid, only phage particles with defective (empty) capsids could be detected. The dimensions of these phages' virions are shown in Table S1.

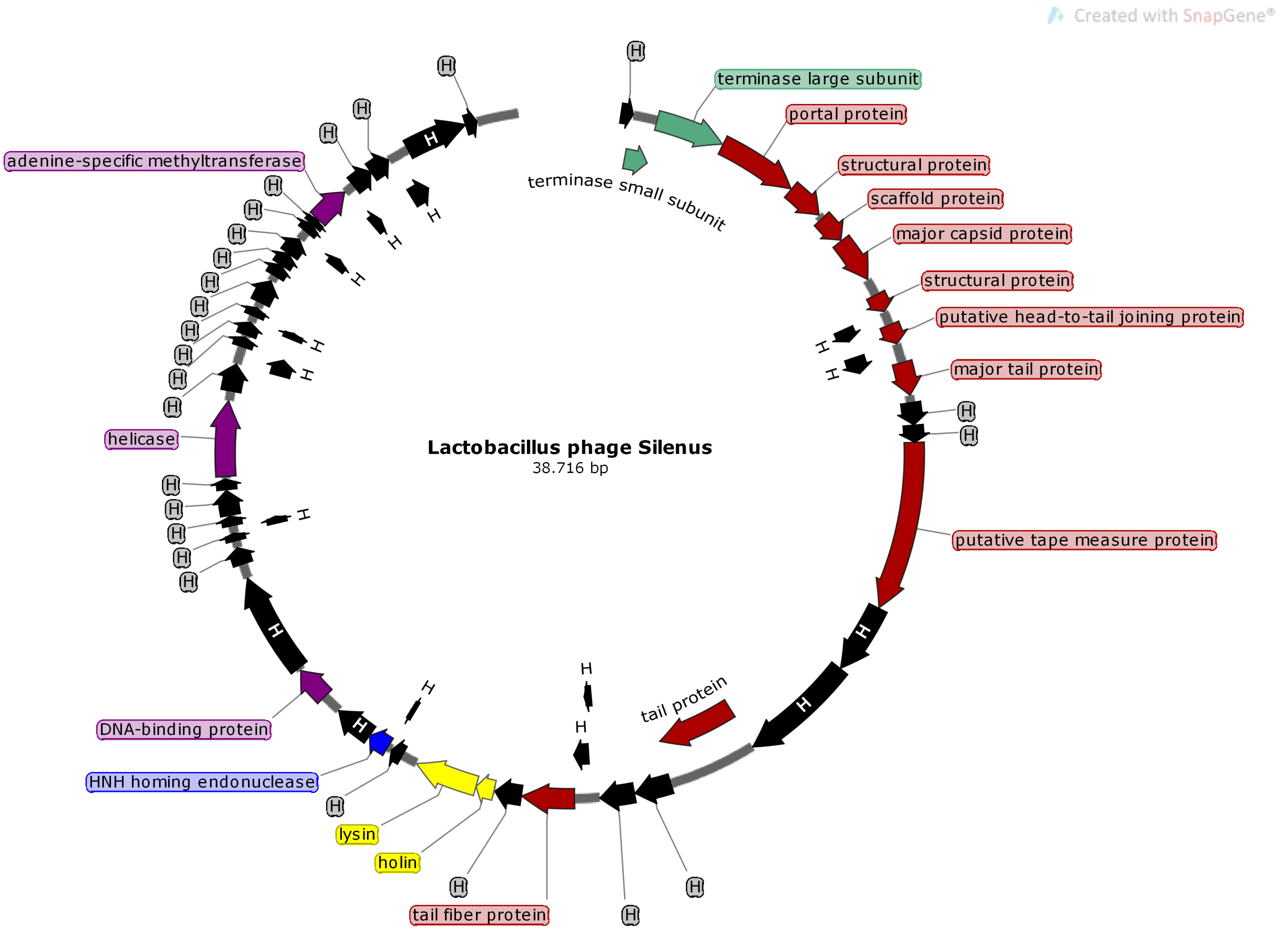


**Figure 3.** Genome map featuring all predicted proteins with assigned functions for phage Silenus. Colour code corresponds to the same protein functions as in Fig. 5.

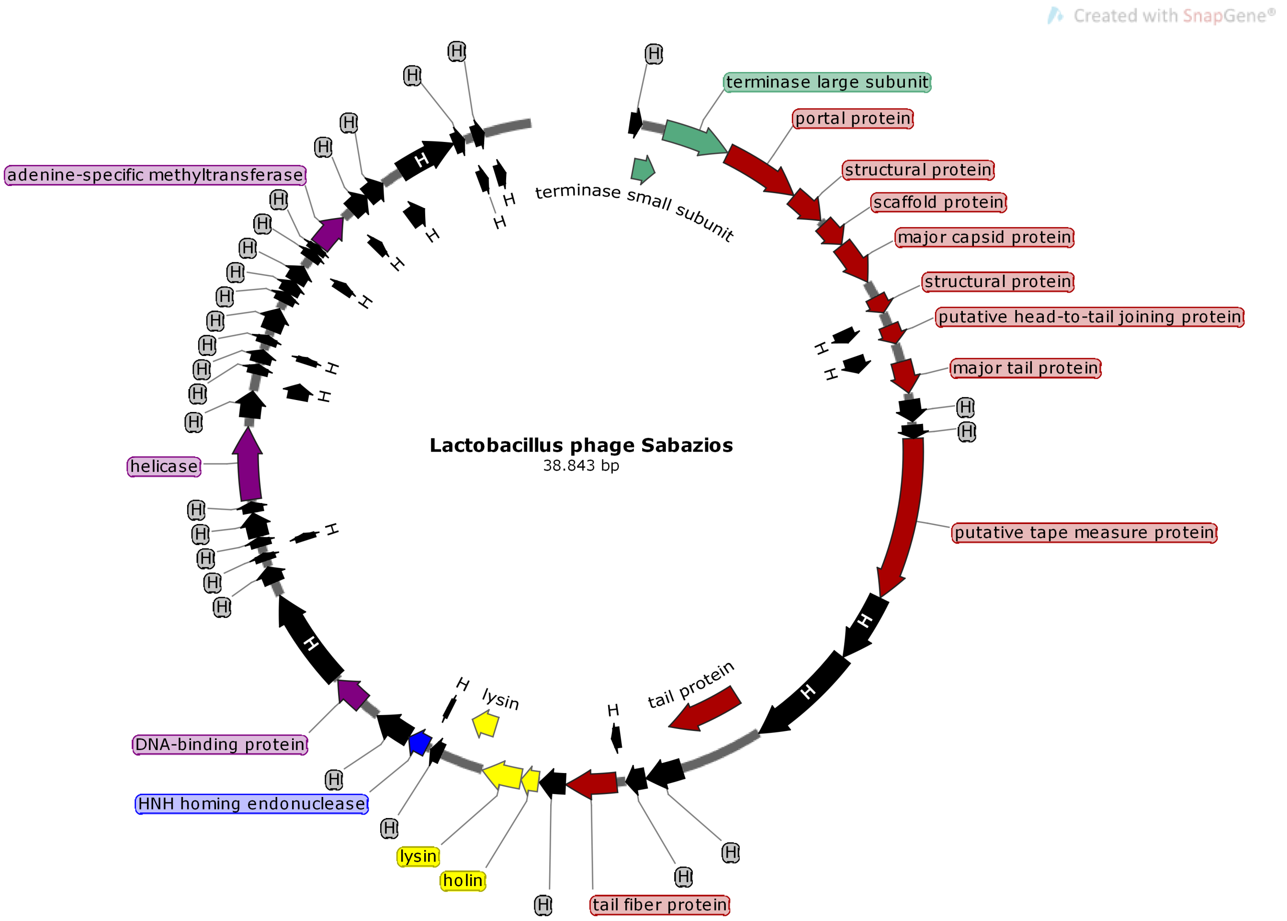


**Figure 4.** Genome map featuring all predicted proteins with assigned functions for phage Sabazios. Colour code corresponds to the same protein functions as in Fig. 5.

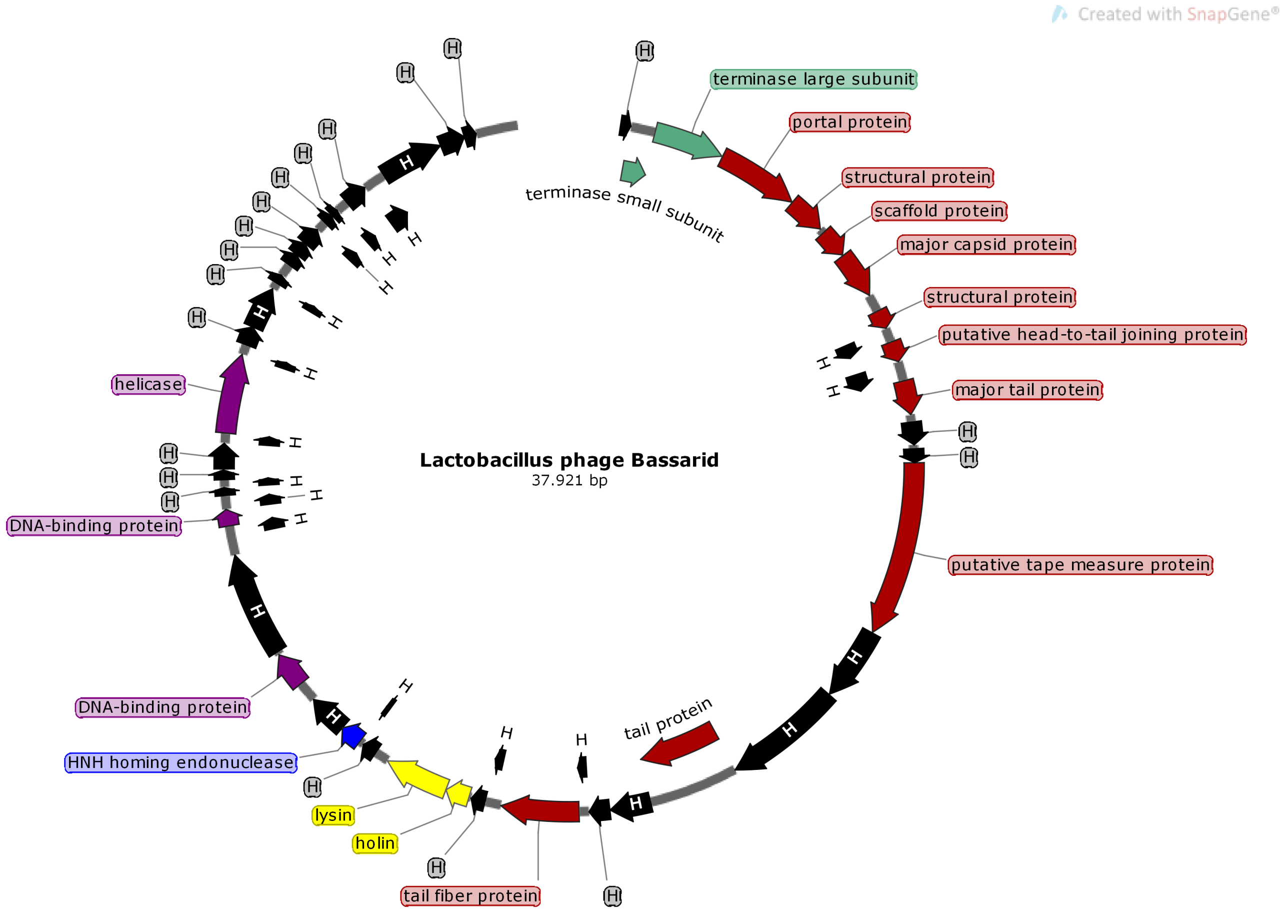


**Figure 5.** Genome map featuring all predicted proteins with assigned functions for phage Bassarid. Colour code corresponds to the same protein functions as in Fig. 5.

```
Reference sequence (1): AUV59728.1
Identities normalised by aligned length.
Colored by: identity

                       cov    pid   1 [         .         .         .         .         :         .         .         . 80
1 AUV59728.1      100.0% 100.0%     MKNKLLLLAASLSLFLMPLTAMASKGDQGVDWAKYQGTNGVFGYSSDKFVISQLGGTVNGSIYEQYTYPTQVASAIAAGK
2 AYH91857.1       64.8%  92.2%     MKKKLLLLLASLALFLMPLTAMASKGDQGVDWAKYQGTNGVFGYSKDKFVISQLGGTVNGSIYEQYTYPTQVASAIAAEK
3 AVH85737.1      100.0%  93.5%     MKKKLLLLLASLALFLMPLTAMASKGDQGVDWAKYQGTNGVFGYSKDKFVISQLGGTVNGSIYEQYTYPTQVASAIAAEK
4 AYH91858.1       46.5%  93.0%     --------------------------------------------------------------------------------
  consensus/100%                    ................................................................................
  consensus/90%                     ................................................................................
  consensus/80%                     ................................................................................
  consensus/70%                     MKpKLLLLhASLuLFLMPLTAMASKGDQGVDWAKYQGTNGVFGYSpDKFVISQLGGTVNGSIYEQYTYPTQVASAIAAtK

                       cov    pid  81         .         1         .         .         .         .         :         . 160
1 AUV59728.1      100.0% 100.0%     RAHTYLWGQFGSSKTQAKAMLDYMLPKVQTPKGSIVALDYEDGASGDKQANTDTIKYALKRIKDFGYTPMLYGYLKYFNA
2 AYH91857.1       64.8%  92.2%     RAHTYLWGQFGSSKAQAKAMLDYMLPKVQTPKGSIVALDYEDGASGDKQANTDAILYALQRIKNAGYTPMLYGYLNYFNA
3 AVH85737.1      100.0%  93.5%     RAHTYLWGQFGSSKAQAKAMLDYMLPKVQTPKGSIVALDYEDGASGDKQANTDAILYALQRIKNAGYTPMLYGYLNYFNA
4 AYH91858.1       46.5%  93.0%     --------------------------------------------------------------------------------
  consensus/100%                    ................................................................................
  consensus/90%                     ................................................................................
  consensus/80%                     ................................................................................
  consensus/70%                     RAHTYLWGQFGSSKsQAKAMLDYMLPKVQTPKGSIVALDYEDGASGDKQANTDsIhYALpRIKshGYTPMLYGYLpYFNA

                       cov    pid 161         .         .         .         2         .         .         .         . 240
1 AUV59728.1      100.0% 100.0%     HVYLSQISGTYKLWLGEYPDYNVTPKPNYNYFPSWENVALFQFTSTYIAGGLDGNIDLTGITDNGYTKNNKPVTNTPAVD
2 AYH91857.1       64.8%  92.2%     HVYLGQISKTYKLWLGEYPDYNVTPKPNYNYFPSWENVALFQFTSTYIAGGLDGNIDLTGITDNGYTKNNKPVTDTPAVD
3 AVH85737.1      100.0%  93.5%     HVYLGQISKTYKLWLGEYPDYNVTPKPNYNYFPSWENVALFQFTSTYIAGGLDGNIDLTGITDNGYTKNNKPVTDTPAVD
4 AYH91858.1       46.5%  93.0%     -----------------------------------------------------MNIDLTGITDNGYTKNNKPVTDTPAVD
  consensus/100%                    .....................................................hNIDLTGITDNGYTKNNKPVTsTPAVD
  consensus/90%                     .....................................................hNIDLTGITDNGYTKNNKPVTsTPAVD
  consensus/80%                     .....................................................hNIDLTGITDNGYTKNNKPVTsTPAVD
  consensus/70%                     HVYLuQIStTYKLWLGEYPDYNVTPKPNYNYFPSWENVALFQFTSTYIAGGLDGNIDLTGITDNGYTKNNKPVTDTPAVD

                       cov    pid 241         :         .         .         .         .         3         .         . 320
1 AUV59728.1      100.0% 100.0%     TGKEAEDTPKADIKKGDKVKVKFSAKHWATGEAIPSWVRGNTYTVTAVSGKKVLLSGINSWIARSNVEILQTNTVQSNVK
2 AYH91857.1       64.8%  92.2%     TGKDADETPKADIKKGEH--------------------------------------------------------------
3 AVH85737.1      100.0%  93.5%     TGKDADETPKADIKKGDKVKVKFSAKHWATGEAIPLWVRGNTYTVTAVSGKKVLLSGINSWIARSNVEILQNKPVQSNVK
4 AYH91858.1       46.5%  93.0%     TGKDADETPKADIKKGDKVKVKFSAKHWATGEAIPLWVRGNTYTVTAVSGKKVLLSGINSWIARSNVEILQNKPVQSNVK
  consensus/100%                    TGK-A--TPKADIKKG-+..............................................................
  consensus/90%                     TGK-A--TPKADIKKG-+..............................................................
  consensus/80%                     TGK-A--TPKADIKKG-+..............................................................
  consensus/70%                     TGKDADETPKADIKKGDKVKVKFSAKHWATGEAIP.WVRGNTYTVTAVSGKKVLLSGINSWIARSNVEILQspsVQSNVK

                       cov    pid 321         .         .         :         .         .         .         .       ] 398
1 AUV59728.1      100.0% 100.0%     LPSGVKRESGTFTANQTLRVWNKPGMSYTGVNYYRGESVRYQGYIRNGNYIYAAYQSSNGTWHYVAVRENGVALGTFK
2 AYH91857.1       64.8%  92.2%     ------------------------------------------------------------------------------
3 AVH85737.1      100.0%  93.5%     LPSGVKREYGTFTANTTLRVWNKPGMAYTGVNYYRGESVRYQGYIRNGNYIYAAYQSSNGAWHYVAVRENGVALGTFK
4 AYH91858.1       46.5%  93.0%     LPSGVKREYGTFTANTTLRVWNKPGMAYTGVNYYRGESVRYQGYIRNGNYIYAAYQSSNGAWHYVAVRENGVALGTFK
  consensus/100%                    ..............................................................................
  consensus/90%                     ..............................................................................
  consensus/80%                     ..............................................................................
  consensus/70%                     LPSGVKRE.GTFTANpTLRVWNKPGMuYTGVNYYRGESVRYQGYIRNGNYIYAAYQSSNGsWHYVAVRENGVALGTFK
```

MView 1.63, Copyright © 1997-2018 Nigel P. Brown

**Figure 6.** Multiple amino acid sequence alignments of the two Sabazios lysins (AYH91857.1 and AYH91858.1, the lysin of Bassarid (AUV59728.1) and the lysin of Silenus (AVH85737.1).
